# Intestinal delta-6-desaturase activity determines host range for *Toxoplasma* sexual reproduction

**DOI:** 10.1101/688580

**Authors:** Bruno Martorelli Di Genova, Sarah K. Wilson, J.P. Dubey, Laura J. Knoll

**Affiliations:** Department of Medical Microbiology and Immunology, University of Wisconsin - Madison, 1550 Linden Drive, Madison, WI 53706, USA; United States Department of Agriculture, Agricultural Research Service, Animal and Natural Resources Institute, Animal Parasitic Diseases Laboratory, Building 1001, Beltsville, MD 20705-2350, USA

## Abstract

Many eukaryotic microbes have complex lifecycles that include both sexual and asexual phases with strict species-specificity. While the asexual cycle of the protistan parasite *Toxoplasma gondii* can occur in any warm-blooded mammal, the sexual cycle is restricted to the feline intestine^1^. The molecular determinants that identify cats as the definitive host for *T. gondii* are unknown. Here, we defined the mechanism of species specificity for *T. gondii* sexual development and break the species barrier to allow the sexual cycle to occur in mice. We determined that *T. gondii* sexual development occurs when cultured feline intestinal epithelial cells are supplemented with linoleic acid. Felines are the only mammals that lack delta-6-desaturase activity in their intestines, which is required for linoleic acid metabolism, resulting in systemic excess of linoleic acid^2, 3^. We found that inhibition of murine delta-6-desaturase and supplementation of their diet with linoleic acid allowed *T. gondii* sexual development in mice. This mechanism of species specificity is the first defined for a parasite sexual cycle. This work highlights how host diet and metabolism shape coevolution with microbes. The key to unlocking the species boundaries for other eukaryotic microbes may also rely on the lipid composition of their environments as we see increasing evidence for the importance of host lipid metabolism during parasitic lifecycles^4, 5^. Pregnant women are advised against handling cat litter as maternal infection with *T. gondii* can be transmitted to the fetus with potentially lethal outcomes. Knowing the molecular components that create a conducive environment for *T. gondii* sexual reproduction will allow for development of therapeutics that prevent shedding of *T. gondii* parasites. Finally, given the current reliance on companion animals to study *T. gondii* sexual development, this work will allow the *T. gondii* field to use of alternative models in future studies.

## Main

The apicomplexan parasite *Toxoplasma gondii* causes a chronic infection in nearly one third of the human population and is well-known for causing congenital infections leading to blindness, mental retardation, and hydrocephaly of the developing fetus. *T. gondii* has a complex lifecycle containing both sexual and asexual phases. The *T. gondii* asexual cycle can occur in any warm-blooded animal when contaminated food or water is consumed and *T. gondii* disseminates throughout the host, converting to an encysted form in muscle and brain tissue. In contrast, the *T. gondii* sexual cycle is restricted to the feline intestinal epithelium, culminating in the excretion of environmentally resistant oocysts^1^. The molecular basis for this species specificity is unknown.

To determine the molecular mechanisms that define the species specificity of *T. gondii* sexual development, we generated cat intestinal organoids (Fig. 1a), then seeded these epithelial cells onto glass coverslips. These monolayers displayed intestinal epithelial properties, including polarization and tight junction formation (Fig. 1b). To simulate natural infection, *T. gondii* was harvested from mouse brains 28-40 days after primary infection and the parasites were released from the brain cysts by pepsin and acid digestion. After neutralization with sodium carbonate, parasites were seeded onto the cat intestinal monolayers, incubated for five days, and stained for markers of the parasite pre-sexual stage called a merozoite^6, 7^. While we observed occasional GRA11B and BRP1 staining, the vast majority of the culture was negative for these merozoite markers (Fig. 1c), suggesting that a required nutrient was limiting under these culture conditions. Because recent studies showed that the *T. gondii* asexual stages scavenge fatty acids, particularly oleic acid, from the host^8^ and that sexual development of many fungi is dependent on linoleic acid^9^, we surmised that supplementation with these fatty acids could facilitate *T. gondii* sexual development. We added 200 μM oleic or linoleic acid to cat intestinal monolayer culture medium 24 hours prior to infection with *T. gondii*. After 5 days of infection, we found that the addition of linoleic acid but not oleic acid caused approximately 35% of the *T. gondii* to express both merozoite stage markers (Fig. 2a). Similarly, GRA11B mRNA was significantly more abundant in cat intestinal cells supplemented with linoleic acid compared to any other condition (Fig. 2b). As seen in vivo cat intestine, GRA11B changes localization from within the parasite dense granule organelles in the early stages of development to the parasitophorous vacuole and parasitophorous vacuole membrane in later stages of development^6^. We see similar localization of GRA11B depending on vacuole size, likely representing early, middle and late stages (Fig. 1e-g). BRP1 has previously been localized to the rhoptry organelles in the apical end of the merozoite^7^, similar to the structures we see in figure 1e-g.

**Fig. 1.**
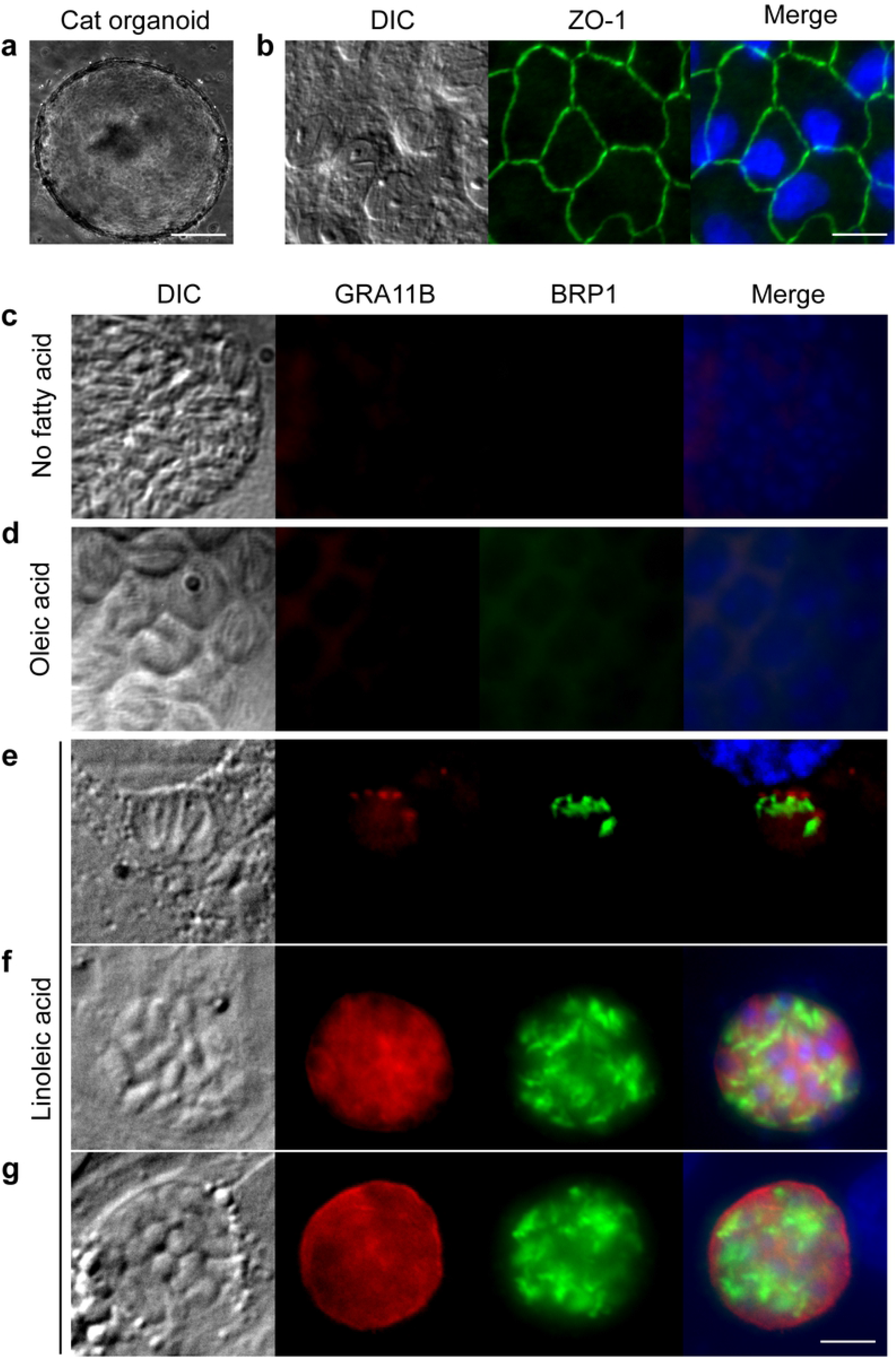
Linoleic acid enhances progression through the sexual stages. **a**, Cat intestinal organoids were generated from small intestine sections and were grown in basement membrane matrix. Example of a growing organoid, 100 μm size bar. **b**, Intestinal organoids were dissociated using trypsin and single cells seeded onto glass coverslips to grow as monolayers. The cells in the monolayer expressed the tight junction protein ZO-1 (green), 20 μm size bar. Cat intestinal monolayers were incubated with either **c**, no fatty acid supplementation, **d**, 200 μM oleic acid, or **e, f, g**, 200 μM linoleic acid for 24 hours, then infected with ME49 bradyzoites for 5 days. Parasites undergoing pre-sexual development were commonly seen only with linoleic acid supplementation as marked by staining with GRA11B (red) or BRP1 (green). Parasites in **e**, early, **f**, middle or **g**, late stages of sexual development were noted by differential localization of GRA11B. All panels are 20 μm square with a 5 μm white size bar in the lower right corner.

**Fig. 2.**
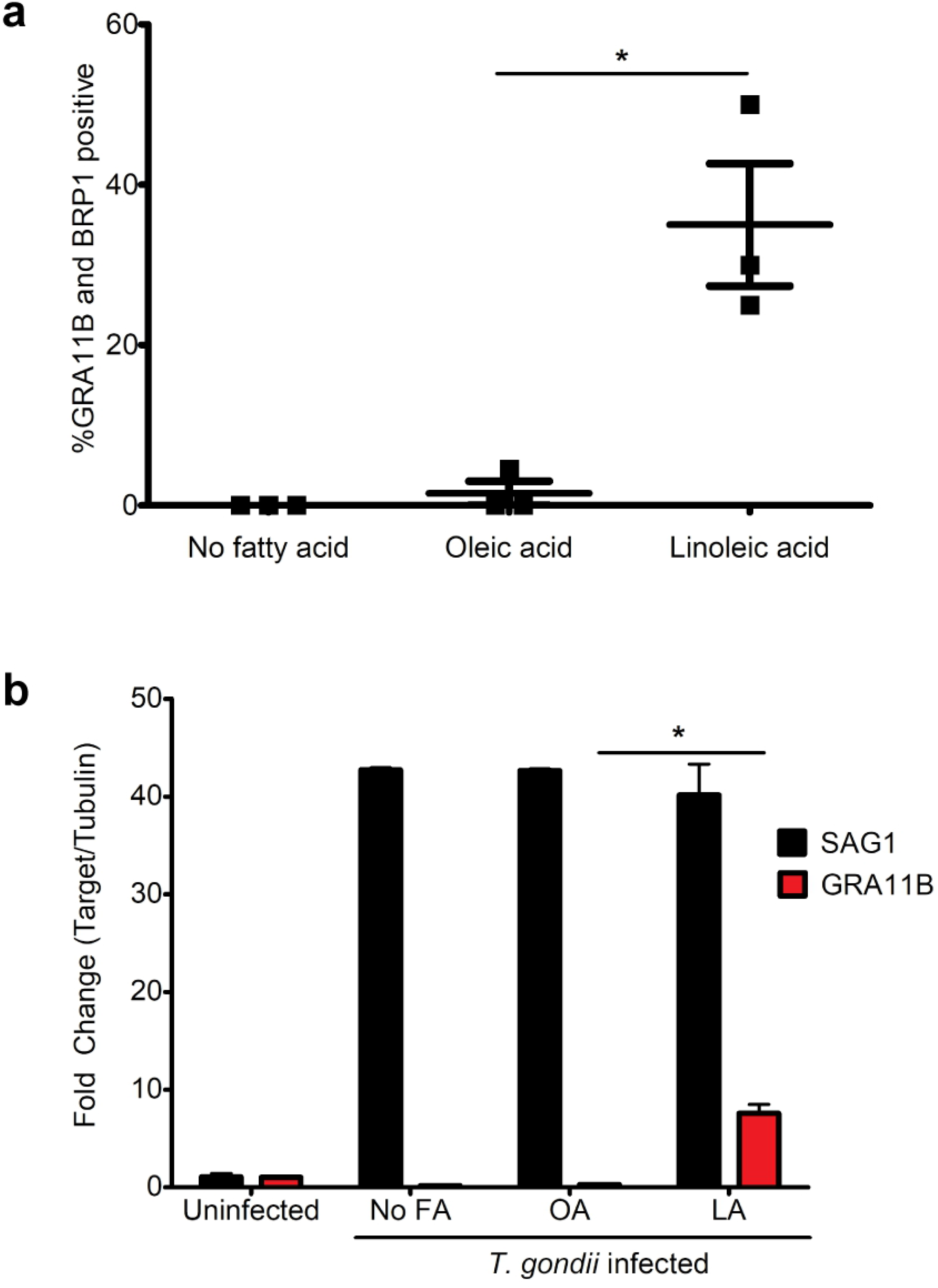
Quantification of merozoites in cat tissue culture. **a**, Cat intestinal organoids were disassociated by trypsin then grown as monolayers on glass slides. Slides were divided into three different groups: not supplemented with fatty acid, supplemented with 200 μM oleic acid or supplemented 200 μM linoleic acid. Monolayers were infected with *T. gondii* ME49 bradyzoites purified from brains of chronic infected mice at a 1:10 MOI. Five days after infection, staining for GRA11B and BRP1 along with DAPI, allowed the percentage of vacuoles positive for GRA11B and BRP1out of the total vacuoles was determined. Total number of parasitophorous vacuoles were counted by positive DAPI staining and confirmed by morphology with DIC. At least 50 parasitophorous vacuoles were counted per replicate. Three biological replicates were counted and on average 35% of the total vacuoles were positive for both GRA11B and BRP1 in the linoleic acid supplemented monolayers. *p-value = 0.0126 with N = 3 by two-tailed unpaired t test. Straining for both BRP1 and GRA11B was used to ensure that merozoite stages were counted. RNAseq and immunofluorescent imaging of the cat intestinal epithelium shows that GRA11B is exclusively expressed in merozoites^6^. BRP1 is a rhoptry protein that was initially found in bradyzoites; however, it is also expressed in merozoites^7^. **b**, Cat intestinal monolayers were grown as described in panel **a**, except monolayers were quenched by TRIzol 5 days postinfection, RNA was extracted, and cDNA was synthesized using an oligo (dT) primer to amplify mRNA. Expression of SAG1 and GRA11B were quantified by qPCR and the fold change calculated in comparison with uninfected cells. TUB1A was used to normalize gene expression across samples. GRA11B expression was significantly more abundant in the linoleic acid supplemented monolayers with two biological replicates. *p-value = 0.0155 with N = 2 by twotailed unpaired t test.

Within the feline intestine, merozoites are known to differentiate into micro- and macrogametes that fuse to become diploid oocysts. After 7 days of infection, we saw round structures with reactivity to the macrogamete protein AO2^10^ in cat intestinal monolayers cultured with 200 μM linoleic acid but not in unsupplemented or oleic acid-supplemented cultures (Fig. 3a-c). PCR of these day 7 linoleic acid supplemented cultures amplified message for AO2 as well as the predicted microgamete flagellar dynein motor protein TGME49_306338 with 44% identity to the homologue from the motile green alga *Chlamydomonas reinhardtii* (Fig. 3d). In parallel, we assessed for the presence of intracellular oocyst wall biogenesis in these linoleic acid supplemented cat cells by using the 3G4 antibody^11^ that recognizes the *T. gondii* oocyst wall. There were approximately 9 oocyst walls per cm^2^ of cultured cat cells with supplemented with 200 μM linoleic acid but none in not supplemented or oleic acid-supplemented cultures (Fig. 3eh). Addition of 20 μM linoleic acid did not enhance oocyst wall production, indicating that the concentration of linoleic acid was critical for proper development.

**Fig. 3.**
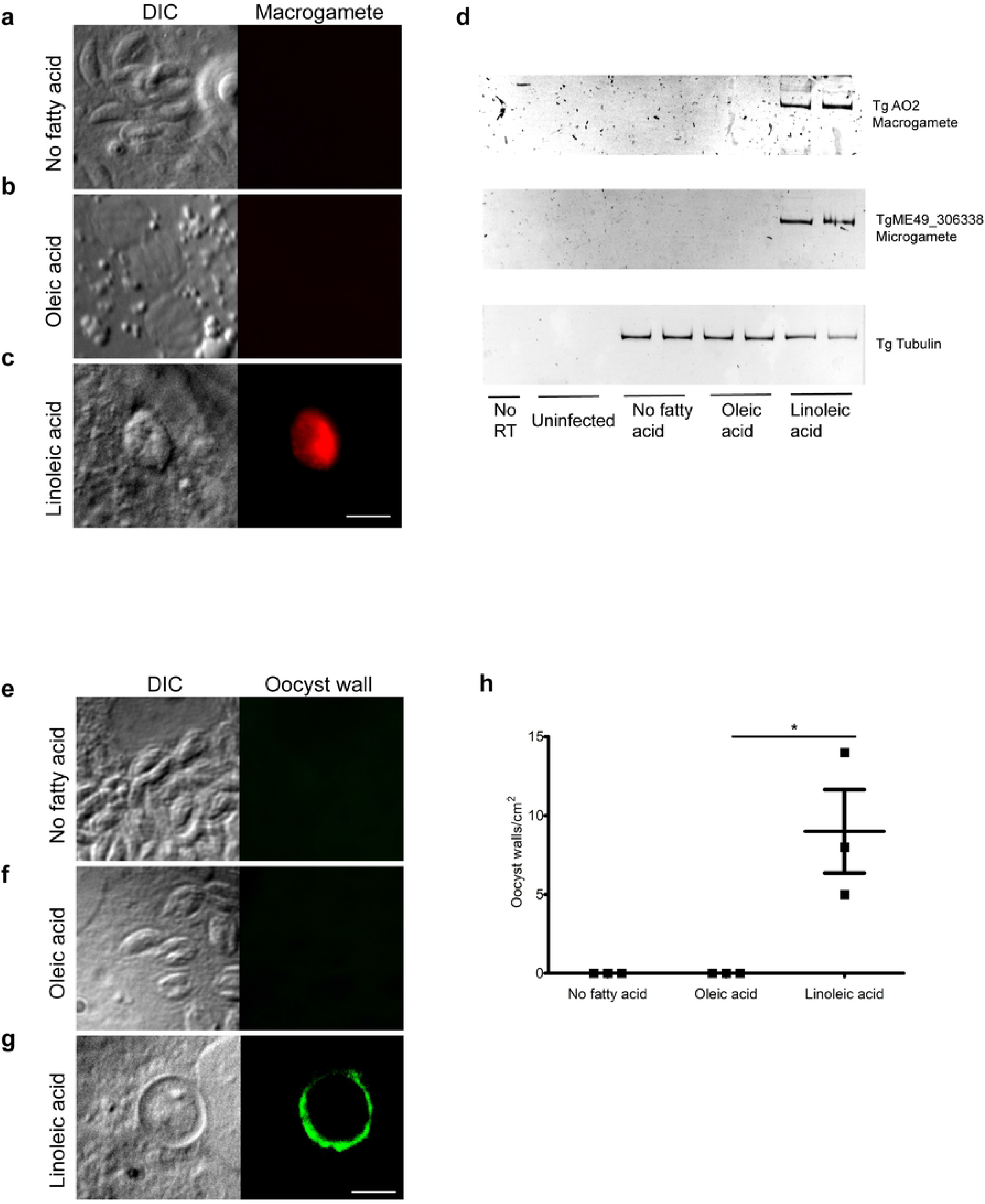
Identification of gametes and intracellular oocysts in cat tissue culture. Cat intestinal organoids were disassociated by trypsin then grown as monolayers on glass slides. Monolayers were grown to confluency and then were incubated with either no fatty acid supplementation (**a** and **e**), 200 μM oleic acid (**b** and **f**), or 200 μM linoleic acid (**c** and **g**) for 24 hours and infected with ME49 bradyzoites purified from brains of chronic infected mice. After 7 days, monolayers were incubated with mouse anti-AO2 (panels **a – c**) or mouse monoclonal IgM 3G4 (panels **e – g**). The amiloride-sensitive amine oxidase, copper-containing protein 2 (AO2) is an enzyme exclusively expressed in macrogametes and early oocysts and has a possible role in oocyst wall biogenesis^10^. AO2 expression was only detected by immunofluorescence in the monolayers supplemented with linoleic acid (panel **c**). 3G4 is a mouse monoclonal antibody produced by immunizing mice with purified oocyst walls^11^, thus it is a marker of oocyst wall biogenesis. Only monolayers supplemented with linoleic acid (**g**) had positive 3G4 vacuoles. All panels are 20 μm square with a 5 μm white size bar in the lower right corner. **d)** Markers for macrogamete and microgamete expression were also evaluated by PCR. Cat intestinal monolayers were grown in 24-well plates until confluency and then infected with *T. gondii* bradyzoites in duplicate using the same conditions as above. Seven days post-infection, RNA was extracted with TRIzol and cDNA was synthesized using an oligo (dT) primer to only amplify mRNA. AO2 was again used as a marker for macrogametes and the expected PCR product is 218bp. To assess microgamete presence, we selected the gene TgME49_306338, which is overexpressed in the gametes stage, corresponded to day 7 post-infection in cats^29^ and has 44% identity to a protein expressed in the flagella of the motile green algae *Chlamydomonas reinhardtii*. The expected PCR product for TgME49_306338 is 160bp. TUB1A was used as an input control and results in a 172bp product. NO RT corresponds to a cDNA synthesis reaction without the addition of reverse transcriptase (RT) as a control for genomic DNA contamination. Equivalent amounts of cDNA per sample were used as a template for each PCR reaction, and the products were separated on an acrylamide gel. Bands with the correct size showing AO2 and TgME49_306338 expression were only observed in linoleic acid supplemented monolayers. **h)** The number of positive oocyst walls stained with 3G4 were quantified. Cat intestinal monolayers were infected with *T. gondii* bradyzoites and after 7 days fixed with 3.7% formaldehyde in PBS and incubated with 3G4 as showed in the panels e, f and g. The number of positive oocyst walls were counted in each slide and divided by the area of slide in cm^2^. The number of positive oocysts walls in monolayers supplemented with linoleic acid was significantly higher than supplementing with oleic acid in three biological replicates. *p-value = 0.0272 with N = 3 by two-tailed unpaired t test.

The dependence of *T. gondii* sexual development on high levels of linoleic acid was intriguing because cats are the only mammal known to lack delta-6-desaturase activity in their small intestines^2, 3^. Delta-6-desaturase is the first and rate-limiting step for the conversion of linoleic acid to arachidonic acid. Linoleic acid is the dominant fatty acid in cat serum, comprising 25-46% of the total fatty acid^12–15^, whereas rodents serum contains only 3-10% linoleic acid^16–19^. We hypothesized that the lack of delta-6-desaturase activity in the cat small intestine allows for a buildup of linoleic acid from the diet, which then acts as a positive signal for *T. gondii* sexual development. To test this hypothesis, we infected mouse intestinal monolayers with *T. gondii* and supplemented them with linoleic acid and the chemical SC26196, a specific inhibitor of the delta-6-desaturase enzyme, to establish high steady-state levels of linoleic acid^20^. Five days after infection of the mouse culture with *T. gondii*, we assessed merozoite markers BRP1^7^ and GRA11B^6^. We detected expression of GRA11B and BRP1 in mouse intestinal cells only when supplemented with both linoleic acid and SC26196 (Fig. 4). These data suggest that the delta-6-desaturase enzyme must be inhibited in order for high enough levels of exogenous linoleic acid to increase and induce *T. gondii* sexual development in non-feline intestinal cells. Similar to cat cells, mouse intestinal monolayers supplemented with both linoleic acid and SC26196 had approximately 26% of the *T. gondii* vacuoles expressing both BRP1 and GRA11B (Fig. S1)

**Fig. 4.**
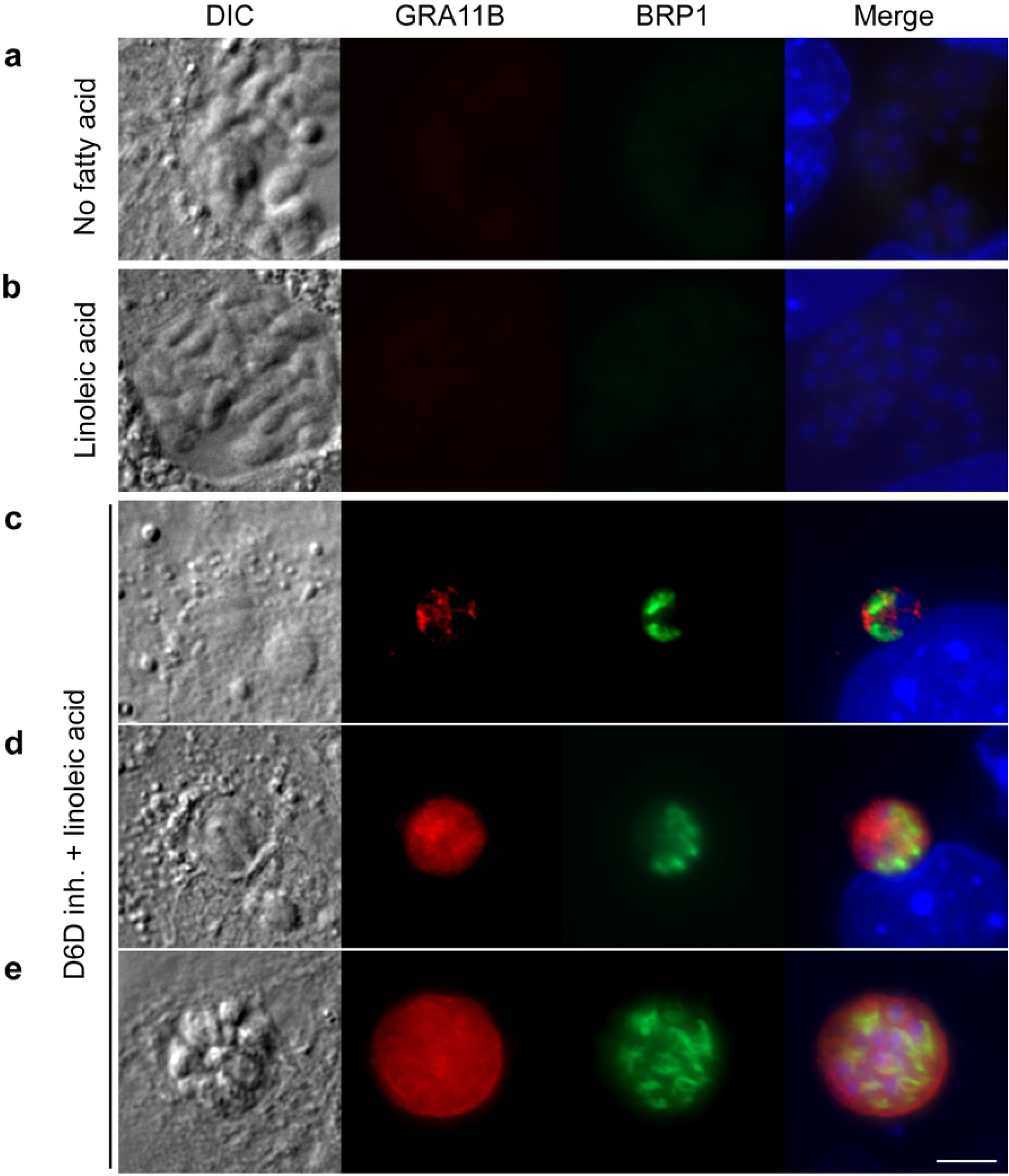
Inhibition of delta-6-desaturase permits sexual development in mouse culture. Mouse intestinal monolayers were incubated with either **a**, no fatty acid supplementation, **b**, 200 μM linoleic acid or **c, d, e**, 200 μM linoleic acid plus the delta-6-desaturase inhibitor SC26196 for 24 hours, then infected with ME49 bradyzoites for 5 days. Only in cultures supplemented with linoleic acid and SC26196 were parasites undergoing pre-sexual development detected by staining with GRA11B (red) or BRP1 (green). Parasites in **c**, early, **d**, middle, or **e**, late stages of development were noted by differential localization of GRA11B. All panels are 20 μm square with a 5 μm white size bar in the lower right corner.

Oocysts excreted in cat feces must undergo a sporulation process to become infectious to the next host. We attempted to sporulate the round structures containing oocyst wall antigen that were derived from either cat or inhibited mouse cultured intestinal cells at room temperature with aerosolization for 7-14 days. Unfortunately, few structures were obtained from the monolayers, they did not appear to sporulate and they were not infectious to mice. We hypothesized that *T. gondii* oocyst development and infectivity would require physiological conditions in a whole animal that could not be recapitulated in tissue culture. To test this hypothesis, we inhibited delta-6-desaturase activity in the intestines of live mice. The delta-6-desaturase inhibitor SC26196 is effective as an anti-inflammatory agent in whole animal experiments^21^. Because it was previously seen that sporozoites shifted to the rapidly replicating asexual stage called a tachyzoite within eight hours after the oral inoculation into rats^22^, we fed the mice a linoleic acid-rich diet and pretreated them with the delta-6-desaturase inhibitor SC26196 (or a no-inhibitor control) 12 hours prior to oral infection with *T. gondii* and every 12 hours thereafter. In mice fed both the linoleic acid-rich diet and the SC26196 inhibitor, seven days after infection, qPCR of ileum cDNA showed high expression of the *T. gondii* merozoite marker GRA11B and low expression of the asexual tachyzoite stage marker SAG1^23^ (Fig. 5a, Table S1). Ileum sections on day seven postinfection were paraffin embedded and stained with hematoxylin & eosin or reticulin stain. Pre-sexual and early oocysts stages were present only in the tissue of mice fed linoleic acid and the delta-6-desaturase inhibitor (Fig. 5b, c).

**Fig. 5.**
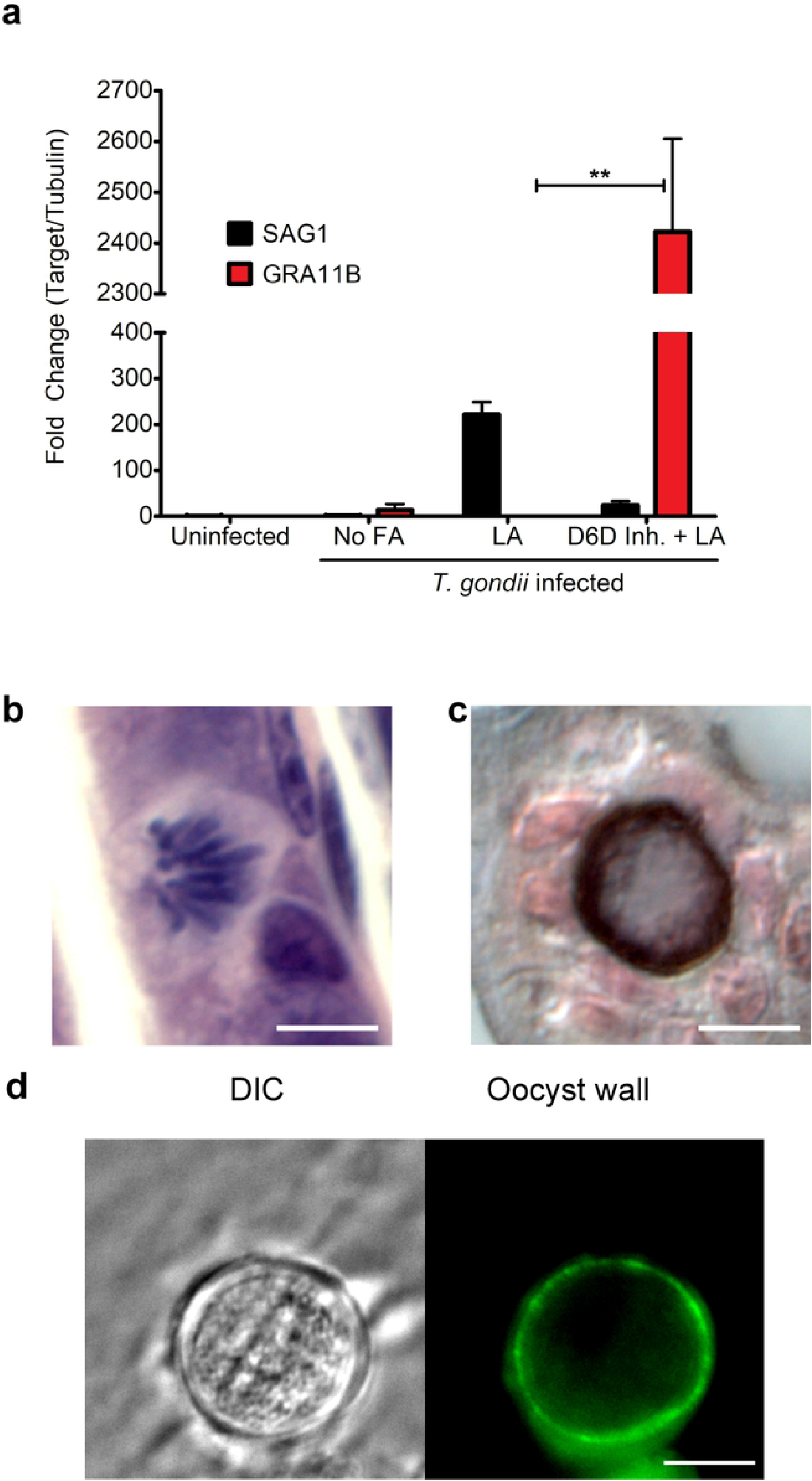
Mice shed oocysts after inhibition of delta-6-desaturase. Mice were gavage fed linoleic acid and the delta-6-desaturase inhibitor SC26196 12 hours prior to infection with ME49 bradyzoites, and then every 12 hours for the 7 days of infection. **a**, qPCR of cDNA from the ileum for tachyzoite marker SAG1 (black) and GRA11B (red) shows that GRA11B is significantly up regulated only in the presence of SC26196 (p-value = 0.0057 with N = 2 by twotailed unpaired t test). **b**, Ileum sections on day seven postinfection were paraffin embedded and stained with hematoxylin & eosin to visualize pre-sexual stages. **c**, early intracellular oocysts were observed in ileums of D6D inhibited mice impregnated with silver (reticulin stain) and photographed using differential contrast imaging to delineate the oocyst wall in dark brown^30^. 10 μm black size bar in the lower right corner. **d**, Fresh oocysts were fixed in 3.7% formaldehyde in suspension, incubated with mouse monoclonal antibody 3G4 overnight, then incubated with goat anti-mouse Alexa fluor 488 secondary antibody. Panels are 20 μm square with a 5 μm white size bar in the lower right corner.

As early as day six postinfection, oocyst-like structures showing 3G4 antibody-positive staining were present in the mouse feces (Fig. 5d) and increased in number until day seven when the mice were sacrificed. qPCR on genomic DNA from mouse fecal samples showed that *T. gondii* genomic DNA was detectable only in mice treated with SC26196 (Fig. 6a, Table S2), indicating that delta-6-desaturase must be inactivated in mice for *T. gondii* sexual stages to develop in the mouse gut. Mice produced 1000-10,000 oocysts/gram dry feces. To increase the number and duration of oocysts shedding, mice were fed the SC2696 inhibitor every 12 hours only until day 5 postinfection. Oocysts were monitored in the feces with the peak of shedding being days 8-9 with between 100,000-150,000 oocysts/gram dry feces (Fig. 6b), which is within the range seen for cats, 2000-1,500,000 oocysts/gram of feces^24, 25^.

**Fig. 6.**
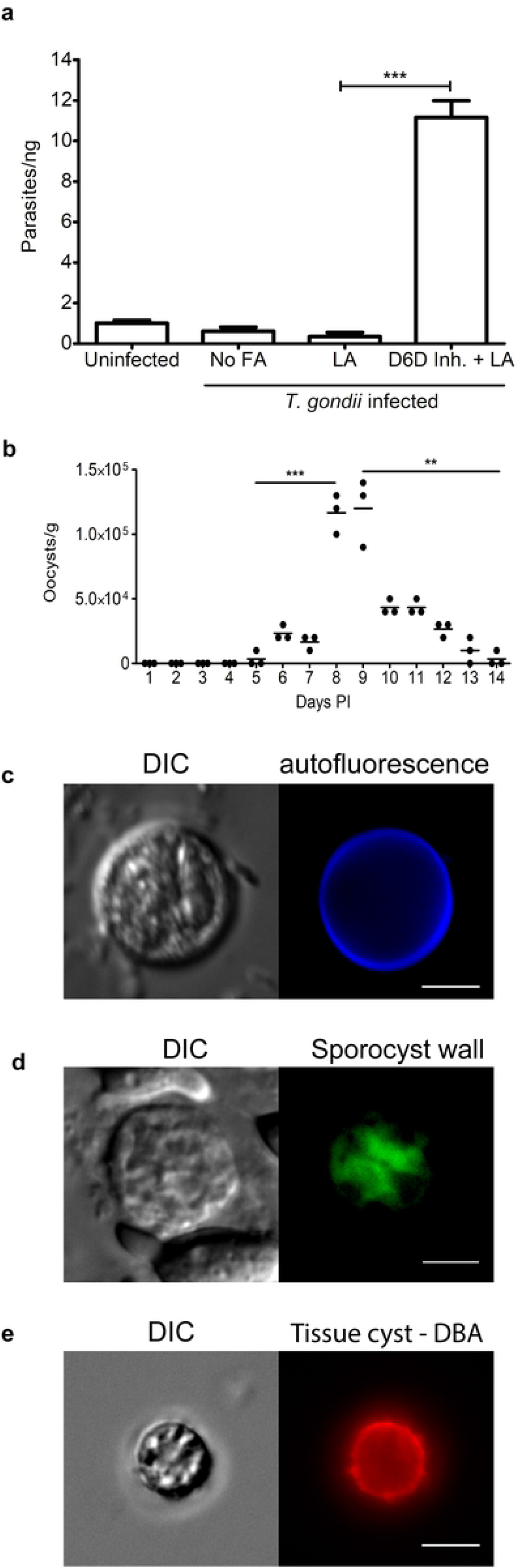
Mouse oocysts are infectious. **a**, qPCR on genomic DNA from mouse fecal samples shows that *T. gondii* genomic DNA is detected only in mice treated with SC26196 (p-value = 0.0002 with N = 3 by two-tailed unpaired t test). **b**, Counts of the number of oocysts/gram of feces over time. ***p-value = 0.0003 day 5 vs 8 and **p-value = 0.0017 day 9 vs 14. **c**, After 7 days in sporulation conditions, sporocysts were visible by DIC, and blue autofluorescence of the oocyst walls was enhanced. All panels are 20 μm square with a 5 μm white size bar in the lower right corner. **d**, To expose the sporocyst wall to the 4B6 antibody^11^, the sporulated oocysts were dried to the slides, then fixed and permeabilized with cold acetone for 30 minutes, incubated with mouse monoclonal antibody 4B6 overnight then incubated with goat anti-mouse Alexa fluor 488 secondary antibody. **e**, At 28 days postinfection with oocysts, *T. gondii* cysts were purified from the brains of mice and detected by *Dolichos biflorus* agglutinin^31^ (DBA, red). All panels are 20 μm square with a 5 μm white size bar in the lower right corner.

*T. gondii* oocysts are susceptible to desiccation, making them unable to sporulate^26^. Therefore the mouse feces or the intestinal contents were immediately placed in saline and sporulated at room temperature with aerosolization. After seven days, sporulation was evident in approximately 50% of the oocysts by visualization of sporozoites, a deep blue autofluorescent wall^27^ (Fig. 6c), and reactivity with the 4B6 antibody that recognizes the two individual sporocysts within the oocysts^11^ (Fig. 6d). The sporulated oocysts were infectious to mice as seen by serum conversion and cysts in the brains 28 days later (Fig. 6e & S2). Similar to oocysts derived from a cat, these mouse-derived sporulated oocysts were stable and infectious for at least three months when stored at 4°C.

All together, these results define the mechanism of species specificity for *T. gondii* sexual development and show that we can break the species barrier for *T. gondii* sexual development by inhibiting delta-6-desaturase activity in the intestines of a non-feline host. The lack of delta-6-desaturase activity and the build-up of linoleic acid likely enhance *T. gondii* sexual development in multiple ways. First, prior work suggests linoleic acid is cytotoxic for the asexual tachyzoite stage^28^, thus tachyzoite development would be halted in a linoleic rich environment. Second, inhibition of delta-6-desaturase likely lowers arachidonic acid levels, which would alter the production of immune lipid mediators known as eicosanoids. Finally, and possibly most important, the dramatic difference between oleic acid with one double bond and linoleic acid with two, highlights that linoleic acid is probably used as a signaling molecule and not to meet basic nutritional needs. Quorum-sensing for sexual reproduction in fungi is dependent on oxygenation of linoleic acid but not oleic acid^9^. The multiple host and *T. gondii* cyclooxygenases and lipoxygenases likely oxygenate linoleic acid to an oxylipin signaling molecule for *T. gondii* sexual development to proceed.

## Acknowledgments

We sincerely thank Jason Spence and his lab for assistance with intestinal organoid culture, Aurélien Dumètre, Adrian Hehl and John Boothroyd for cat stage specific antibodies. Maria Arendt for assistance with intestinal pathology images. Heather Fritz, David Ferguson and Jean François Dubremetz for advice. Christina Hull, Benjamin Rosenthal and Rodney Welch for editing of the manuscript.

## Author Contributions

BMGD conducted all of the organoid culture experiments, SKW performed all of the qPCR, BMDG and LJK conducted all of the mouse experiments. JPD provided the ME49 *T. gondii* oocysts for mouse infections. Experimental design was conducted by BMDG with assistance from SKW, JPD and LJK. BMDG and LJK wrote the manuscript. BMDG, SKW, JPD and LJK reviewed and edited the manuscript.

## Methods

### Ethics Statement

Mice were treated in compliance with the guidelines set by the Institutional Animal Care and Use Committee (IACUC) of the University of Wisconsin School of Medicine and Public Health (protocol #M005217). Cats were treated in compliance with the guidelines set by the IACUC of the United States Department of Agriculture, Beltsville Area (protocol #15-017). Both institutions adhere to the regulations and guidelines set by the National Research Council.

### Intestinal organoids

Cat intestinal organoids were established from jejunum sections obtained from fetal small intestinal sections. Mouse intestinal organoids were established from jejunum sections from 8-week-old C57BL/6J male mice. Organoids were generated as described previously^32^. Briefly, intestinal sections were washed in ice cold PBS containing 0.1 mg/mL streptomycin and 100 U/mL penicillin for 20 minutes. Sequentially, EDTA (Sigma) was added to a final concentration of 2 mM and the tissue incubated for 40 minutes at 4°C. The tissue was then rinsed in cold PBS without EDTA and vigorously shaken until crypts were seen in the supernatant. The crypt suspension was filtered using a 70 μm cell strainer and the crypts were centrifuged at 80 x g for 5 minutes. The cells were resuspended in Matrigel (BD Biosciences), pipetted into a 24 well plate, allowed to polymerize and then covered with organoid medium. The organoid medium contains Advanced DMEM/F12 with 2 mM Glutamax, 20 mM HEPES, 1 x B27, 1 x N2, 10% v/v Fetal bovine serum, 10 mg/L Insulin, 5.5 mg/L Transferrin, 0.67 mg/L Selenite, Penicillin and Streptomycin (all from Invitrogen), 50 ng/ml human EGF (R&D systems), 10 mM Nicotinamide (Sigma), 3 μM CHIR99021 and 10 μM Y-27632 (both Selleckchem) and 50% v/v conditioned medium obtained from L-WRN cell line (ATCC CRL 3276). The medium was changed every other day and the organoids were expanded by passing the cells through a 25 gauge needle every week. All experiments were done with cells at passage 2 to 5 and cells were regularly checked for mycoplasma contamination (MicoAlert Lonza).

### Intestinal monolayers and fatty acid supplementation

Monolayers were generated from intestinal organoids as described previously^33^. Briefly, established cat or mouse intestinal organoids were washed with cold PBS, digested by 0.05 % m/v trypsin for 5 min at 37 °C, centrifuged at 250 x g for 3 minutes and resuspended in fresh pre-warmed organoid medium. Cell suspension was added into a chamber slide (Thermo) pre-coated with Entactin-Collagen IV-Laminin (Corning) for cat cells or 2% m/v Gelatin in PBS (Sigma) for mouse cells. The slides were coated by air drying the basement membrane matrix or gelatin to air dry overnight. The monolayers were grown for 10-15 days prior to infection with *T. gondii* bradyzoites, with media change every other day until cells reached 90% or more confluency. Linoleic acid or oleic acid conjugated to BSA (Sigma) was added to the organoid monolayers to 0.2 mM 24 hours prior to infection.

### Bradyzoite preparation and infection

C57BL/6J mice were oral gavage infected with 500-1000 ME49 oocysts from cat feces. At 28 days postinfection, brains were removed, washed in cold PBS and homogenized with a glass tissue grinder. The suspension was centrifuged at 400 x g for 10 minutes and the pellet suspended in 20% m/v Dextran (Average MW 150,000, Sigma). Bradyzoite cysts were pelleted and separated from brain material by centrifugation at 2200 x g for 10 minutes. The pellet was washed in PBS, digested by 0.1 mg/mL pepsin in HCl for 5 minutes at 37°, then neutralized with an equal volume 1% Sodium Carbonate (Sigma). Bradyzoites were spun at 250 x g for 10 minutes, resuspended in pre-warmed organoid medium and added onto the organoid monolayers with a multiplicity of infection of 1 bradyzoite: 10 intestinal epithelial cells (MOI 1:10).

### Delta-6-desaturase inhibition

SC 26196 (Cayman) was solubilized in DMSO and used at 20 μM in mouse organoid monolayers. For *in vivo* treatment, the inhibitor was solubilized in 0.5% m/v methylcellulose and the mice were given 50 mg/kg every 12 hours by oral gavage^21^. 4-week-old C57BL/6J female mice deleted in Z-DNA-binding protein^34^ were divided into four different groups: uninfected control, *T. gondii*-infected without fatty acid supplementation, *T. gondii*-infected with linoleic acid supplementation, and *T. gondii*-infected with linoleic acid and SC 26196 inhibitor. Each mouse supplemented with linoleic acid received 10 μL of 99% linoleic acid oil (MilliporaSigma Cat#843483) suspended in 0.5% Methylcellulose per day by oral gavage. Mice were infected with 1000 brain cysts purified as described above by oral gavage and euthanized 7 days post infection. Sample size was at least 2 mice per group and the experiment was repeated 5 times. Alternatively, each mouse was infected with one mouse brain at least 2 months postinfection with at least 1000 cysts. Mice were treated with SC 26196 until day 5 postinfection. Feces were collected from days 5-14 and oocysts enumerated by microscopy.

### Immunofluorescence

Intestinal organoid monolayers or mouse fecal samples were fixed in 3.7% formaldehyde in PBS for 20 minutes, permeabilized with 0.2% triton X-100 (Sigma) in PBS at room temperature for one hour and then blocked with 3% BSA in PBS at room temperature for one hour. Primary antibody was incubated at 4°C overnight in 0.2% v/v Triton x-100 and 3% BSA in PBS (1:100 mouse anti-GRA11B, 1:100 rabbit anti-BRP1, 1:100 mouse anti-AO2, 1:50 monoclonal mouse anti-ZO1 (Santa Cruz) or 1:25 mouse IgM anti-oocyst wall 3G4. Sporulated oocysts from mouse feces were dried onto slides, fixed and permeabilized with ice cold acetone for 30 minutes and incubated with 1:20 mouse 4B6 to the visualize the sporocyst. Slides were incubated one hour with the specific secondary antibody (1:500 goat antirabbit Alexa Fluor 488 and 1:500 goat anti-mouse Alexa Fluor 594) at room temperature for one hour and then washed 3 times with PBS. Cells nuclei were stained with 10 μM DAPI (Sigma). Slides were mounted in Vectashield antifade mounting medium (VectorLabs). Samples were imaged on Zeiss Axioplan III equipped with a triple-pass (DAPI/fluorescein isothiocyanate [FITC]/Texas Red) emission cube, differential interference contrast optics, and a monochromatic Axiocam camera operated by Zen software (Zeiss) and processed using ImageJ (Fiji packet).

### Tissue sectioning and histology

Ileums were fixed in 3.7% formaldehyde in PBS overnight, embedded in paraffin and sectioned by the Translational Research Initiatives in Pathology laboratory at the University of Wisconsin-Madison. The sections were stained with hematoxylin & eosin or reticulin (Silver) stain.

### Real-time PCR on ileum cDNA

Mice with and without delta-6-desaturase inhibitor treatment were euthanized 7 days post infection. The ileum of each mouse was removed and homogenized in 1mL of TRIzol. Total RNA was isolated according to manufacturer’s protocol (Invitrogen) and treated with amplification grade Dnase I. cDNA was generated using the Invitrogen SuperScript III First-Strand Synthesis kit with random hexamer primers. GRA11B and SAG1 were used as markers of sexual and asexual stages, respectively. The *T. gondii* housekeeping gene TUB1A was used to normalize target gene expression. Real-time quantitative PCR was performed using Bio-Rad iTaq Universal SYBR Green Supermix on an Applied Biosystems StepOnePlus Real-Time PCR system. The efficiency of each primer set was calculated from the slope of a 1:10 dilution standard curve of tachyzoite gDNA, where E = 10^(−1/slope). The Pfaffl method^35^, which accounts for differences in efficiencies, was then used to calculate the relative gene expression of GRA11B and SAG1 per sample, in triplicate. Only wells with one melt curve temperature were used, indicating a single product. Primer sequences were as follows:

TUB1A Forward: 5’ – GACGACGCCTTCAACACCTTCTTT – 3’
Reverse: 5’ – AGTTGTTCGCAGCATCCTCTTTCC – 3’
SAG1 Forward: 5’ – TGCCCAGCGGGTACTACAAG – 3’
Reverse: 5’ – TGCCGTGTCGAGACTAGCAG – 3’
GRA11B Forward: 5’ – ATCAAGTCGCACGAGACGCC – 3’
Reverse: 5’ – AGCGAATTGCGTTCCCTGCT – 3’

### Real-time PCR for *T. gondii* genomic DNA in fecal samples

Fecal samples from the mice with and without delta-6-desaturase inhibitor treatment were collected. Genomic DNA was generated from 0.1 grams of feces from each mouse using the power soil DNA kit (QIAGEN) according to the manufacturer’s instructions except that cells were broken by a bead beater instead of a vortex. A standard curve was generated using a dilution series of 10^1^–10^5^ parasites per well amplified using the SAG1 primer set described above, based on a genomic DNA sample with known parasite quantity. The Ct values were plotted against the log of the parasite numbers. The number of target gene copies in each sample can be interpolated from the linear regression of the standard curve.

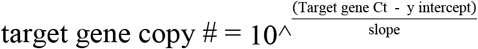

Real-time PCR was performed on each sample, in triplicate, using Bio-Rad iTaq Universal SYBR Green Supermix on an Applied Biosystems StepOnePlus Real-Time PCR system. The calculated copy numbers of each sample were normalized based on the ng of nucleic acid used as PCR template. Only wells with one melt curve temperature were used, indicating a single product.

### PCR of cat intestinal monolayers

Cat intestinal monolayers were grown in 24-well plates until confluency and then were incubated with either no fatty acid supplementation, 200 μM oleic acid, or 200 μM linoleic acid for 24 hours. The monolayers were infected with ME49 bradyzoites purified from brains of chronic infected mice in duplicate with uninfected monolayers as a negative control. 7 days post-infection, RNA was extracted with TRIzol and cDNA was synthesized as described above. TgAO2 was used as a marker for macrogametes and TgME49_306338 was used as a marker for microgametes. TUB1A was used as an input control using the same primers as above. A cDNA synthesis reaction without the addition of reverse transcriptase was used as a control for genomic DNA contamination. Equivalent amounts of cDNA per sample were used as a template for each PCR reaction, and the products were separated on an acrylamide gel and imaged. Primer sequences were as follows:

TgAO2 Forward: 5’ – GTCTTGGTTCGTTGAAGGGGCTG – 3’
Reverse: 5’ – CGTCCTCGATGCCCATGAAATCTG – 3’
TgME49_306338 Forward: 5’ – CCACGTCCTTCGCCGATG – 3’
Reverse: 5’ – CATCAGAGGTCCCAGGTTGTCG – 3’

### Statistical methods

All real-time PCR fecal samples were run in triplicate technical replicates. The difference between the mean target gene copy numbers was analyzed by two-tailed unpaired t tests. The real-time PCR intestinal samples were run in triplicate from two biological replicates per group. The difference between the mean relative expression of each target gene was analyzed by two-tailed unpaired t tests.

### Oocyst sporulation and mouse infections

Fresh fecal samples were obtained from each mouse, homogenized in PBS and then centrifuged at 1500 x g. The pellet was resuspended in PBS plus penicillin and streptomycin and the samples were shaken for 7 to 14 days at room temperature in presence of oxygen. Mice oocysts were stable for at least 3 months at 4°C. Naïve mice were infected with approximately 250 mouse oocysts through intraperitoneal injection. Mice were humanely euthanized at day 28 postinfection, their brains removed, homogenized and either incubated with biotinylated DBA 1:1000 or purified with 20% m/v Dextran as described above before DBA incubation. All cysts were then incubated with streptavidin-conjugated AlexaFluor-594 1:1000 and imaged on Zeiss Axioplan III equipped with a triple-pass (DAPI/fluorescein isothiocyanate [FITC]/Texas Red) emission cube, differential interference contrast optics, and a monochromatic Axiocam camera operated by Zen software (Zeiss) and processed using ImageJ (Fiji packet).

